# The spatiotemporal richness of hummingbird wing deformations

**DOI:** 10.1101/2023.05.08.539717

**Authors:** Dimitri A. Skandalis, Vikram B. Baliga, Benjamin Goller, Douglas A. Altshuler

**Affiliations:** Department of Zoology, University of British Columbia; Psychological and Brain Sciences & Johns Hopkins Kavli Neuroscience Discovery Institute, Johns Hopkins University; College of Agriculture Data Services, Purdue University

**Keywords:** shape, flight, plasticity, dynamics

## Abstract

Animals exhibit an abundant diversity of forms, and this diversity is even more evident when considering animals that can change shape on demand. The evolution of flexibility contributes to aspects of performance from propulsive efficiency to environmental navigation. It is, however, challenging to quantify and compare body parts that, by their nature, dynamically vary in shape over many time scales. Commonly, body configurations are tracked by labelled markers and quantified parametrically through conventional measures of size and shape (descriptor approach) or non-parametrically through data-driven analyses that broadly capture spatiotemporal deformation patterns (shape variable approach). We developed a weightless marker tracking technique and combined these analytic approaches to study wing morphological flexibility in hoverfeeding Anna’s hummingbirds (*Calypte anna*). Four shape variables explained >95% of typical stroke cycle wing shape variation and were broadly correlated with specific conventional descriptors like wing twist and area. Moreover, shape variables decomposed wing deformations into pairs of in- and out-of-plane components at integer multiples of the stroke frequency. This property allowed us to identify spatiotemporal deformation profiles characteristic of hoverfeeding with experimentally imposed kinematic constraints, including through shape variables explaining <10% of typical shape variation. Hoverfeeding in front of a visual barrier restricted stroke amplitude and elicited increased stroke frequencies together with in- and out-of-plane deformations throughout the stroke cycle. Lifting submaximal loads increased stroke amplitudes at similar stroke frequencies together with prominent in-plane deformations during the upstroke and pronation. Our study highlights how spatially and temporally distinct changes in wing shape can contribute to agile fluidic locomotion.

**Summary:** Hummingbirds exhibit complex wing deformations throughout the stroke cycle, and the timing and origin of these deformations differs between hoverfeeding behaviours.

## INTRODUCTION

A moving animal’s instantaneous pose, its configuration, is computationally represented by tracking landmarks on the body, curves representing its outline, or reconstructed surfaces of the body itself (Cheney et al., 2020; Ellington, 1984; Hedrick, 2008). Shape is the geometric information remaining after factoring out differences in scaling, translation, and rotation amongst all configurations (Dryden and Mardia, 2016; Kendall, 1984; Slice, 2001). Behaving animals, particularly with deforming bodies, are challenging to quantify because the same landmark information dictates the analysis of both shape and kinematics (Yezzi and Soatto, 2003). Choices in the design of the landmark set, including distribution, density, and tracking error, affect both the instantaneous configuration and its alignment to other configurations. Landmarks are often chosen according to visibility and anatomical significance, including joints and wing tips, but the sparsity of natural features can cause a low-pass filtering of the geometry.

The dynamics of moving animals are therefore often tracked by augmenting feature sets with (potentially high-variance) points like the estimated shoulder location (e.g., Altshuler et al., 2012; Read et al., 2016; Tobalske et al., 2007). Alternatively, physical markers may be applied, provided they do not interfere with the body dynamics (Bender et al., 2015; Hedrick et al., 2012; Iriarte-Diaz et al., 2012; Riskin et al., 2008; Song et al., 2014). Additional decisions may be required, such as the choice of measurements derived from the tracked configurations or the choice of metric when comparing configurations directly. A major question is therefore how different methods relate to each other, and how they influence our inferences on animal movements.

Given a body quantified by its instantaneous configurations, there are different methods to identify changes in motion and shape. We distinguish between parametric *descriptors* of a configuration’s size and shape comprising pre-selected measurements, and data-driven approaches to comparing configurations directly to derive new, non-parametric *shape variables*.

Parametric descriptors of a configuration’s size and shape commonly include dimensional measurements like length and area, measurements of shape as quantified by ratios of dimensions and internal angles, and bespoke indices like tip roundedness (e.g., Baliga et al., 2019; Lockwood et al., 1998; Lucas et al., 2014; Matloff et al., 2020; Riskin et al., 2008; Tobalske et al., 2007; Zheng et al., 2013).

Parametric descriptors simplify comparisons among individuals, species, and studies because the measurements should be independent of the details of the configuration or shape spaces. Conflicting terminology and definitions nonetheless persist (Stiles and Altshuler, 2004). Some non-dimensional morphological parameters are well-studied in engineering contexts and have proven useful for understanding morphological flexibility in behaving animals. Wing twist, the change in angle from tip to base, is an essential factor in determining the propulsive efficiency of rotating and flapping wings (Ingersoll et al., 2018; Leishman, 2000; Zheng et al., 2013). Camber, the maximum height deviation of a propulsor section, allows lifting surfaces to generate greater weight-supporting forces at lower flight speeds; both birds and bats have been observed to rely on cambering in slow flight (Muijres et al., 2008; Muijres et al., 2012). Unfortunately, it is not straightforward to map the wing dimensions of posed specimens to the complex postures of freely behaving animals (Riskin et al., 2010). Propulsor size and shape can vary by species (Riskin et al., 2010) or even among individuals between behaviours. Individual bats provide an extreme example of divergent combinations of adopted kinematics and wing shapes across flight speeds and during force loading (Iriarte-Diaz et al., 2012). It may be difficult to choose a set of descriptors that capture the dynamics of highly flexible and variable body parts’ natural shapes.

Selecting a descriptor set is further challenged when different body shapes appear only during specific behaviours. In that case, without studying the entire range of behaviours, we have only a partial knowledge of the natural extent of deformations. In general, any such set of descriptors will also be spatiotemporally correlated, like serial camber sections through a wing, or causally related, like twist-induced camber (Ennos, 1988). These issues present statistical challenges for identifying the timings and sources of dynamic shape variation among behaviours. They may also introduce bias when crucial elements of shape changes are not included.

Non-parametric approaches to studying body shape involve direct comparisons among configurations (Adams and Cerney, 2007; Dryden and Mardia, 2016; Kendall, 1984). Rather than defining functions on the configurations (such as calculating area or twist), a metric for the similarity between configurations is chosen, such as the Procrustes distance among labelled landmarks (Dryden and Mardia, 2016; Kendall, 1984; Slice, 2001). Common transformations among configurations are factored out, and new shape variables are derived from the major axes of variation through decomposition approaches like singular value decomposition (or, generally equivalently, principal components analysis; Dryden and Mardia, 2016; Schlager, 2017). Projecting configurations onto this new basis enables visual and statistical comparisons of the spatiotemporal deformation signatures of different behaviours. Although efficient, the shape variables may be difficult to interpret beyond the sample set and in real terms. In practice, the number of shape variables (preserved components) is usually chosen statistically or heuristically to achieve a desired compression ratio (Schlager, 2017; Zheng et al., 2013). The biological significance of very low variance components may also not be apparent. On the other hand, like when choosing a descriptor set, pre-selecting the degree of retained information in this way risks losing information about characteristic shape differences and their timings among behaviours.

We study wing shape variation in Anna’s hummingbird (*Calypte anna*) as a model of morphological flexibility during active behaviours. Hummingbirds are spectacularly agile and can generate burst forces several times their body weight through kinematic modulation. Whether dynamic wing shape variation also contributes to maneuvering flight is poorly described (Ravi et al., 2020; Read et al., 2016; Segre et al., 2015; Skandalis et al., 2017). In part, this may be because hummingbirds appear to exhibit much less shape variation than birds with distally morphing wings (large span ratios; Baliga et al., 2019; Riskin et al., 2012; Stowers et al., 2017; Tobalske et al., 2007). However, at the high wing beat frequencies of hummingbirds, the timing of shape and dimensional changes may be as important as their magnitude. We therefore examined wing shape over consecutive stroke cycles as hummingbirds were challenged to perform kinematically constrained hoverfeeding in front of a visual mask and during submaximum load lifting. Hummingbirds adapt their stroke kinematics to these challenges, particularly through greatly reduced or increased stroke amplitudes respectively (Mahalingam and Welch, 2013; Wells, 1993). We anticipate that understanding the relationships between parametric and non-parametric approaches to shape analysis will lead to new insights into the timing and sources of morphological variation among distinct behaviours. Overall, our work demonstrates how the spatiotemporal profiles of wing deformations can contribute to hummingbirds’ performance envelopes.

## METHODS

Our work is organized as follows. (1) We describe our method for tracking large numbers of wing markers through multiple stroke cycles across individuals. We then detail our methods for inferring spatiotemporal deformation patterns through conventional size and shape descriptors together with well-known techniques in biological shape analysis (geometric morphometrics). (2) We review the challenges and our practical approaches to comparing shape variation among individuals when the marking schemes cannot be guaranteed to be similar. (3) For each individual in the data set, we establish the correspondence between parametric and non-parametric measurements of size and shape. The non-parametric shape variables are constructed through singular value decomposition of aligned wing configurations. (4) We study all individuals together to identify spatiotemporal deformation patterns that distinguish typical from kinematically constrained hoverfeeding.

### Wing and body marking

All procedures were approved by the University of British Columbia Animal Care Committee in accordance with the guidelines set out by the Canadian Council on Animal Care. Five adult male *Calypte anna* were wild caught on the campus of the University of British Columbia and housed in an animal care facility. Marker placement dictates what aspects of shape variation can be quantified, such as joint positions (Hedrick et al., 2012; Riskin et al., 2008), the wing perimeter (Ingersoll et al., 2018; Song et al., 2014), fin ray and feather rachis deflection (Bozkurttas et al., 2009; Maeda et al., 2017; Standen and Lauder, 2005), or marker- and markerless reconstructions of internal surfaces (Bachmann et al., 2012; Cheney et al., 2020; Cheney et al., 2022; Deetjen et al., 2017; Gillies et al., 2011; Heinold and Kähler, 2015; Walker et al., 2009; Windes et al., 2018). Most hummingbird wings are featureless and have few landmarks, unlike the veins and patterns on insect wings (Koehler et al., 2012; Le et al., 2013; Phan et al., 2017; Young et al., 2009) and so three-dimensional reconstructions necessitate dense application of markers over the entire wing (Windes et al., 2018). For our desired marker density, painting (e.g., titanium dioxide, (Song et al., 2014)) introduces handling stress, mechanically disrupts preening and wing integrity by interfering with the interlocking barbules (Matloff et al., 2020), and adds inertia, all of which may substantially disrupt aerodynamics and sensory feedback to alter typical flight patterns. Instead, we devised a weightless marker technique (the ‘hummingbird salon’) that yielded long-lasting and weightless markers and afforded substantial time for the animal to recover prior to experiments. We bleached spots on the surface of the wing with a commercially available hair bleach foaming agent, L’Oréal Paris® Perfect Blondissima Crème™ (because they’re worth it).

Awake birds were gently restrained in a tensor bandage with a slot for one wing and laid in a foam cradle with the wing extended through gentle pressure on the outer primaries. We applied at least three dots of bleaching foam to each feather, generally along the rachis of the feather, on selected dorsal covert feathers, and on the leading edge up to the estimated shoulder location. The sixth secondary flight feather (S6) was absent or infrequently visible among birds, and so excluded. The bleaching agent was allowed to dry for at least five minutes, and then thoroughly soaked and rinsed with multiple washes of tap water to ensure that bleaching agent residue could not be ingested by a preening bird. We did not observe any difference in health or flight capacity between marked and unmarked birds in our population. The entire marker set was applied over multiple sessions of about 30 minutes to limit handling stress to the bird. Both marking experience and the small size of the animal relative to the restraining hands led to some differences in the number and placement of markers, resulting in 58-64 markers over the whole wing. This marking technique is not practical on the body feathers because of low contrast and large feather displacement compared to flight feathers. Instead, we applied five white paint spots (Bic® Wite-Out or Paper Mate® Liquid Paper) to the bird’s back on the day of experimentation. The mass of these spots is minor compared to the body inertia (about 0.07 mg).

### Marker tracking

Markers were manually tracked from at least four colour high-speed camera views (one Miro4 and three Miro120s, Vision Research, Inc.), filming at 2200 Hz and 512×512 pixel resolution. Depending on availability, we also used a greyscale Phantom v12.1 and Phantom v311 (Vision Research, Inc.).

Camera frame synchronisation was controlled by a function generator (Tektronix AFG3021B), and shutter time was a maximum of 150 μs, depending on the camera’s capabilities and lens aperture. We used 24-, 32-, and 50-mm lenses, and due to the sensor crop factor and small image size in the center of the lens, nonlinear image distortions were negligible. Cameras were calibrated using direct linear transform coefficients (Hedrick, 2008) determined by a physical calibration object or through a custom calibration wand and sparse bundle adjustment (Theriault et al., 2014). The gravity vector was determined from the suspended physical calibration object’s +Z axis or by dropping a bead through the cameras’ fields of view. Feather motions and variation in lighting through the stroke cycle sometimes resulted in ambiguous marker positions. Where possible, the marker position was estimated based on nearby markers, or for short gaps of a few frames, the position was interpolated by a smoothing spline.

Otherwise, the marker for that individual was deleted from the dataset. Markers at the wing base (secondary flight feathers) were most affected but are densely packed, limiting the impact on reconstruction. The digitisation precision of each marker in each frame was estimated by the reprojection error of the reconstructed 3D position. We enforced a minimum digitising root mean square error (RMSE) of 1. The marker standard deviation was calculated by bootstrapping (100 iterations) the residual error in each camera view and recalculating the 3D reconstructed position (Hedrick, 2008). Smoothed marker positions were computed by a cubic spline weighted by the relative standard deviations of all points in the time series. This fitting procedure resulted in the smoothest function consistent with the marker positions and the digitising error. The smoothing method is implemented in a custom Matlab package (Hedrick, 2008).

Overall, the appearance of spatiotemporal features of marker motions depends on the quality of evidence (e.g., number of camera views on marker, precision of calibration, digitisation precision). High frequency features, such as rapid displacements, require more evidence and denser spatial and temporal sampling. Enforcing a minimum digitising error therefore amounts to a low-pass filter dependent on the magnification of the marker in each camera view. This is because the pixel error will translate to a larger uncertainty for less magnified markers, and so sharp positional changes without strong evidence will be smoothed out. In terms of shape reconstruction, our labelling and tracking method is consequently conservative (type I error) and may miss some real features (type II error).

### Hovering flight challenges

The experimental flight chamber was made of five panels of clear acrylic measuring 52 cm on the sides and top. The bottom panel was covered by a thin 1 cm^2^ mesh to prevent recirculation of the downwash. The tip of a custom feeder was centrally placed 18 cm and >3 wing lengths from the wall (based on average *C. anna* wing length 5.5 cm). The feeder mouth was extended with a tube ∼1.5 cm to force the bird to adopt a similar head position at each entry, although some twisting of the body still occurred. Each bird was provided with a perch on a mass balance (Ohaus Scout Pro) and allowed to acclimate to the chamber for at least one day before experimentation. We used operant conditioning to train the bird to associate activation of filming lights and the experimenter placing the feeder in the chamber with the start of feeding bouts. To minimise the ON duration of the four 500W halogen bulbs, we required feeding to begin within 30 s, or the session was skipped (rare among well-trained birds). In general, filming time was <30 s, and the feeder was immediately removed. Body weight during the bout was recorded as the average of pre- and post-feeding measurements. This filming period is shorter than the length of time for a hummingbird feeding to satiation, so we immediately inspected the filming, repeating the session if required. Otherwise, the bird was allowed to feed until it returned to the perch while the recordings were downloaded for 10-15 minutes, when the feeder was removed again. In addition, after every three trials the bird was allowed to feed ad libitum for a further 15 minutes.

Hummingbirds were recorded flying either with unconstrained postures (unchallenged ‘typical’ wing configurations), while challenged to hoverfeed in front of a barrier placed around the mouth of the feeder (‘visual mask’), or while performing submaximum load lifting. When hoverfeeding near a visual obstacle, such as a large flower or this mask, hummingbirds avoid wing collisions with the obstacle by greatly reducing stroke amplitude and increasing stroke frequency and stroke plane angle (Wells, 1993). We constructed a thin plastic mask with the majority of the interior cut out. The side length of the mask was 5.75 cm, about 55% of the wingspan. The remaining thin border was nearly parallel with the wing direction of motion, so we expected minimal aerodynamic interference, though this cannot be discounted. We did not observe any collision between the wing and the mask. Feeding duration was a few seconds compared to 10 seconds or longer at the unmasked feeder, and consisted of briefly darting into the feeder, sometimes with their necks extended. We also challenged birds through submaximum load lifting trials (Mahalingam and Welch, 2013; Wells, 1993), because aerial agility in hummingbirds is strongly correlated with the capacity to generate large sustained and burst forces (Altshuler et al., 2010a; Segre et al., 2015; Sholtis et al., 2015). When hoverfeeding with weights, hummingbirds increase stroke amplitude but changes in stroke frequency are minimal and irregular (Mahalingam and Welch, 2013). Birds were fitted with beads on an elastic band looped around the neck, equal to 20-25% of body weight, depending on the birds’ cooperation and any evidence of flight distress. One individual refused to hover normally with any added weight.

### Wing anatomy

We dissected the left wing of a male Anna’s hummingbird (*Calypte anna*) euthanised in an unrelated experiment to understand feathers’ functional groupings in terms of actuating elements. The musculoskeletal system and feather insertions were exposed and photographed in a spread posture on a custom light table. Positions of the bones, feathers, and connective tissue between the feathers were traced in Adobe Illustrator. The anatomy was verified in two additional dissections.

### Kinematics and mesh construction

We define the wing position in terms of its azimuth, elevation, and the geometric angle *α* of the least squares plane passing through the centroid of the labelled set. The elevation angle and *α* are respectively reported with respect to the horizontal and vertical planes of the lab reference frame. Stroke reversal timings and duration were determined from the position peaks and were used to find the stroke frequency. The mid-downstroke and mid-upstroke timings were similarly determined as half the downstroke and upstroke intervals, respectively. We constructed a mesh representation of the wing from the labelled points by interpolating forty points along the leading and trailing edges of the mid-downstroke pose, joining spanwise paired points to construct chordal line segments (Song et al., 2014). Internal points were spaced at approximately 0.5 mm and connected through Delaunay triangulation, pruning faces with centroids outside the wing perimeter (2D mesh). We then used the custom Matlab function *gridfit* (D’Errico, 2017) to find a mesh scheme fit to the internal wing markers and with a smooth gradient in all directions. To obtain a consistent 3D mesh, we selected the mid-downstroke as the template frame and then projected it onto the mesh at each time step using the *iso2mesh* Matlab toolbox (Tran et al., 2020). In general, the resulting meshes visually fit the data well although we noted some oversmoothing of real, high-curvature features of the secondary flight feathers and wing base during the upstroke. Animations of the tracked landmarks and wing reconstructions are included as Supplementary Materials.

The scaling of force relationships in flapping wings has been well studied, including aspects of wing kinematics, size, and shape that require special attention (Ellington, 1984; Kruyt et al., 2014; Skandalis et al., 2017). The force equation *F* = *qSC_F_* relates forces *F* to the dynamic pressure *q*, the wing planform area *S*, and a dimensionless force coefficient *C_F_*. Given some assumptions about the dynamics of force production, the dynamic pressure can be resolved into 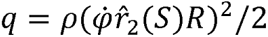, the product of air density ρ, angular velocity 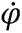, and the non-dimensional second moment of wing area, 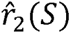, and wing length *R* (Ellington, 1984). Wing area was determined as the surface area of the mesh. Area moments of posed specimens are often calculated through the product of a section blade radius and chord width (Ellington, 1984; Kruyt et al., 2014). Because the mesh has area elements *dA*, we quantify the squared second moment of area at every time slice by 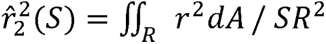, where *r*, *R*, and *S* are respectively the radius of the centre of each element from the wing base, the wing length as above, and the 3D mesh area. Wing twist, camber, and spanwise bending are prominent contributors to variation in *C_F_* throughout the stroke cycle. We approximated the wing twist from tip to root *α* as Θ = *α_tip_* - *α_root_* (see Ingersoll et al. (2018) for an alternative calculation based on the average twist rate between sections). The chordwise camber at each line segment on the wing was determined by *h*_chord_/*c*, where *h*_chord_ is the maximum height of the chordwise strip above its line segment of length *c* (Ennos, 1988). Spanwise camber was computed by a line connecting the wing tip to a point lying at 0.05*c* behind the shoulder projected onto the 3D mesh, which yielded the wing length *R*, mesh height *h*_span_, and finally spanwise camber as *h*_span_/*R*. Individual morphological measurements are reported in Supplementary Table 1 together with posed wing measurements for two individuals, calculated from wing silhouettes as usual (Ellington, 1984).

### Decomposition of shape variation

Two shapes are considered equal if, through scaling, translation, and rotation, they can be superimposed with arbitrary precision. We give some basic details of this approach and point the reader to more detailed discussions elsewhere (Dryden and Mardia, 2016; Guigui et al., 2021; Slice, 2001) and to the implementation in the R package *Morpho* (Schlager, 2017). Shape variation throughout the stroke cycle for each individual was quantified with respect to a template configuration with *k* tracked landmarks *X_M_* ∊ ℝ^3k^. Configurations at times *t*, X*_t_*, were aligned through least squares to the template as X_t_ = λ (X_M_ + d)Γ+ b, with a scaling factor λ ∊ ℝ^+^, special orthogonal rotation Γ ∊ SO (3), and translation b ∊ ℝ^3^ of the configuration centroid. The matrix d ∊ ℝ^3*k*^ contains the time-specific residuals from the template, i.e., the elements of shape, that forms the squared Procrustes distance from the template Δ^2^ = tr(*dd^T^*), which is related to the Riemannian distance by *ρ* = 2 sin^-1^(Δ/2) (Slice, 2001). The template was recovered after iteratively registering all wing configurations until convergence, and so does not correspond to a particular time slice. Factoring out transformations, together with this metric, induces a non-linear shape space, and so some coordinate linearisation is necessary to apply linear statistics. We use an orthogonal projection into the tangent space of the template (a function of the inverse exponential map; Dryden and Mardia, 2016; Schlager, 2017; Slice, 2001), yielding tangent plane coordinates (Dryden and Mardia, 2016; Kendall, 1984; Slice, 2001), *T*_M_*X*. The tangent space coordinates were then decomposed through the (economy-size) singular value decomposition, (*T_M_X*)*_m_* = *UD_m_V^T^*, where the columns of *UD_m_* ∊ ℝ^*t*×3*k*^ are the projection onto the new basis formed by *V* ∊ ℝ^3*k*×3*k*^. (Decomposing the transposed data matrix exchanges the senses of *U* and *V*, as for example in Riskin et al., 2008.) The proportion of variance explained by each of the ranked singular values *d* ∊ *D* is given by *d*^2^/ ∑*d*^2^. Following this procedure, we refer to *UD_m_* as shape scores and *V* as shape variables. The shape variation represented in the *m*th shape variable was determined by zeroing out all but the *m*th singular value and then visualising the reconstructed shape for different shape scores (Dryden and Mardia, 2016). The reconstructed configurations were used to study the orientations of the vectors in ℝ^3^ from the template landmarks to the landmarks of the configurations recovered from the shape scores. The average angle of these vectors to the *xy* plane (π/2 – φ, where φ is the vector’s elevation in spherical coordinates) was used to measure the contributions of in- and out-of-plane deformations to each shape variable. We used custom Matlab scripts to find the frequencies of peak power/frequency in each shape variable.

### Appropriateness of shape analysis methods

Some difficulties arise when applying and tracking landmarks among individuals and behaviours. First, marker placement on hummingbirds’ small and visually homogeneous wings is challenging. As a result, our marker placements and numbers were unequal, violating the assumptions of shape analyses based on labelled landmarks. As a practical solution, we observe that as the number of applied markers increases, the data increasingly resemble a point cloud (under some regularity conditions like isotropic densities and reparametrisation invariance). A point cloud represents the technical limit of reconstructed shape information, and therefore of the latent shapes that we aim to track with our labelled set. We cannot tell when exactly such a density has been reached, so we instead propose the weaker condition that the spatiotemporal information represented in the marker set should appear to be saturating. To investigate this condition, we used Watanabe’s landmark sampling curves to compare the sums of squares obtained by fitting subsets of landmarks to the complete set in each individual (Watanabe, 2018). With sufficient marker density, we expect the sampling curve to plateau, implying that little or no new shape information is gained by adding more markers. The main impact of this assumption, further incorporated into our statistical modelling as detailed below, is that we cannot discriminate individual variation from heterogeneity in landmark placement. Our second consideration is that as a matter of sampling, some hummingbirds did not sustain hovering in every condition. The experimental design is therefore unbalanced, which could influence the decomposition of the shape variance. We therefore constructed the shape variables from the typical hoverfeeding trials only and then projected the wing configurations in each treatment onto this basis. The shape variation we examined is therefore in terms of typical, unchallenged hoverfeeding. We expect this to be the most representative of hummingbird flight in general and not dependent on specific details of the experimental conditions.

Our ultimate population comparison of shape differences among flight conditions relies on the equivalence of the quantified shape information. We examined the consistency of the derived shape variables on an individual-by-individual basis. First, we examined whether the shape variables appeared to contain saturating shape information, as detailed above. Second, we examined whether the shape variables pertained to similar deformations, including the composition of in- and out-of-plane components. Third, we studied the temporal consistency of the shape scores. Finally, we examined whether the shape variables exhibited similar temporal relationships and linear dependencies on the descriptors, as described below.

### Statistical analyses

All statistical models, except where indicated, were fit in the R package *brms* (Bürkner, 2017) with four chains of 2000 iterations each. We calculated Bayesian model *R*^2^ as a measure of the model goodness of fit (Bürkner, 2017; Gelman et al., 2019). Where appropriate, we determine statistical significance based on whether the 95% credible interval excludes 0.

Stroke-averaged kinematics were compared among treatments by calculating the stroke frequency (as 1/stroke duration), amplitude, and plane angle for each stroke cycle. We analysed all three parameters together as a function of flight condition to account for their mutual correlations, and with individual stroke cycles nested within individual bird. This model was regularised through a horsehoe prior with two degrees of freedom (Bürkner, 2017; Carvalho et al., 2009).

Our initial step in examining the relationships between shape variables and descriptors was to examine their correlations. For time series, cross-correlations provide information on both the magnitude and time lag of relationships. This is expressed as ρ_score,variable_ = corr(score, descriptor*_t-k_*) between the shape scores and each kinematic or descriptor variable for time lags *t-k*. The time lags are reported in fractions of the stroke cycle. For this analysis, lags with correlation significantly greater than zero were identified through robust *t*-statistics as implemented in the R package *testcorr* (Dalla et al., 2022). For the reasons discussed above, this was done separately for each individual, resulting in small differences in time lag for each individual. The lags were therefore rounded to the closest 0.02 phase and the number of significant (*p*<0.05) coefficients summed. We show the lags for which all relationships in that period were significant. This ad hoc test was principally used to determine whether the relationships between shape variables and descriptors were similar among individuals.

In principal components analysis, variable loadings are formed by projecting the variables back onto the derived components. Building on this idea, we regressed the shape descriptors with the shape variables. All measurements were standardised so that the effect of the coefficients was interpretable on the same scale (β). For this regression analysis, we did not include temporal autocorrelations, as our goal was to model how the shape scores reflect instantaneous rather than time-lagged associations. Responses were modelled with respect to wing azimuthal position and elevation, wing twist, area, 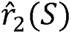, and the chord- and span-wise cambers. The kinematic parameters were included to account for the possibility of mutual correlations between the shape variables and descriptors to stroke phase rather than to each other. We included per-bird random slopes to account for potential inconsistencies in the relationships among birds with differing marking schemes or interindividual variation in morphing patterns. All models were fit with flat priors for all coefficients in *brms*.

Variation in flight behaviour could ostensibly arise through wing shape changes at any interval of the stroke cycle. To compare intervals in which shape significantly differed across behaviours among individuals, we normalised each stroke cycle to unit duration. This resulted in two or three repeated measures (tracked stroke cycles) for each bird in each treatment. We used statistical splines to construct a population reference smooth function that captured the temporal variation in the shape scores during typical hoverfeeding flights (*mgcv*: Wood (2017); p-spline with adaptive and cubic cyclic basis). Differences between treatments were then examined through parametric and non-parametric terms (not to be confused with the parametric and non-parametric approaches to analysing wing size and shape). The parametric term encapsulated average differences in shape score over the whole stroke cycle. The non-parametric term captured the time series trends of the shape scores and the temporal intervals in which each treatment differed from the reference function (smooth difference spline, Δsmooth; p-spline with cubic cyclic basis). Random wiggliness in the smooth functions encompassed interindividual variation in the shape variable time series arising from biological or technical sources (factor-smooths with continuous first derivatives; Wood, 2017). We accounted for autocorrelation because we sought to identify population differences between behaviours, and the shape score at any temporal interval will depend on the stroke cycle history (i.e., lagged effects caused by stroke cycle sampling variation). The autocorrelation coefficient for each shape variable was determined by the value that minimised Akaike’s information criterion (0.95, 0.94, 0.91, and 0.88 on the first to fourth shape variables, respectively). All analyses were fit through restricted maximum likelihood implemented in the R package *mgcv* (Wood, 2017) and smoothed estimates and confidence intervals were constructed with the package *gratia* (Simpson, 2023). The average fits for each treatment were obtained by adding the Δsmooths to the reference function and zeroing out the factor-smooths.

## RESULTS

### Tracking kinematic and shape variation throughout the stroke cycle

Wing configurations tracked by landmark coordinates result from a combination of kinematics and changes in wing shape (Figure 1, S1 Video). In typical hoverfeeding (Figures 1, 2 A), stroke amplitudes averaged 129° (95% CI: 121–139°) with a stroke frequency of 41 Hz (38–44 Hz) and a stroke plane angle of 13.3° (95% CI: 9.1–17.6°). When hoverfeeding in front of a visual mask (Figures 1, 2 A) designed to limit the stroke width, amplitude decreased by 46° (95% CI: 41–51°) and stroke frequency concomitantly increased by 13 Hz (95% CI: 11–15 Hz) and stroke plane angle by 4.2° (95% CI: 0.4–7.8°). Hoverfeeding during submaximum load lifting induced nearly the opposite kinematics (Figures 1, 2 A), with an increase in stroke amplitude of 19° (95% CI: 13–25°) and a decrease in stroke plane angle by 6° (95% CI: 1.5–10.5°), though without a significant decrease in stroke frequency (−1 Hz, 95% CI: −3–1 Hz).

**Figure 1.**
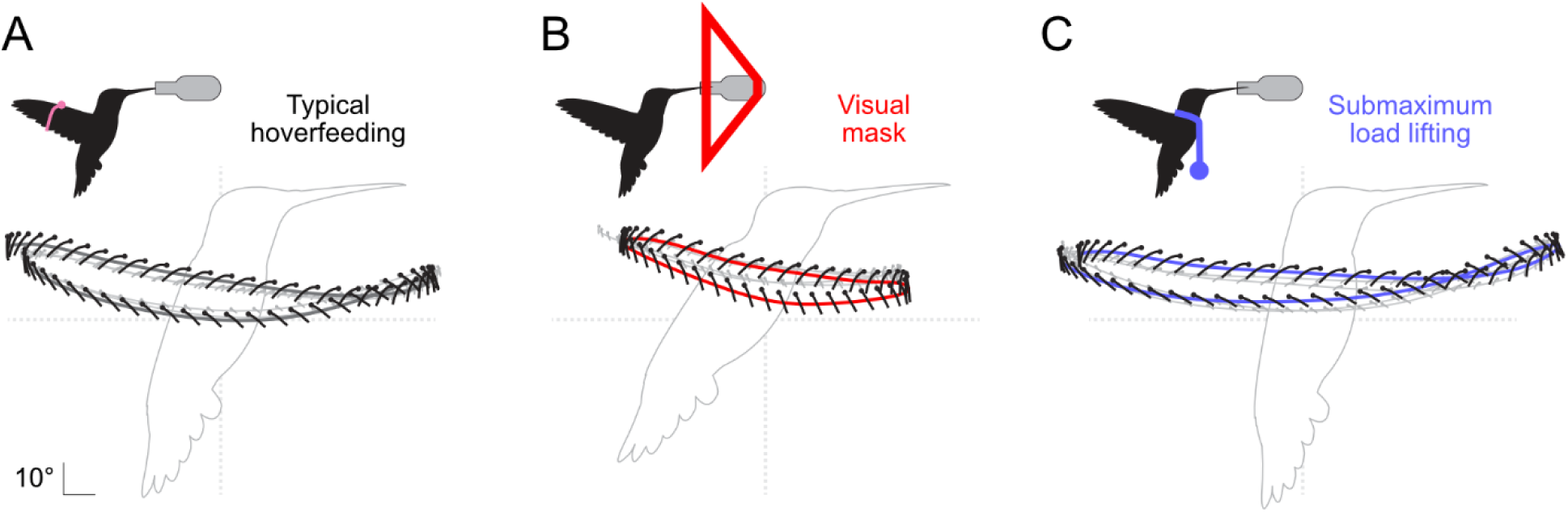
Exemplar hummingbird stroke cycles during typical and kinematically constrained hoverfeeding conditions. One stroke cycle is represented by a thick trajectory line and others in grey. The midspan wing angle and curvature are depicted by a ball-and-stick model on a normalised chord length. Body angle was estimated from body markers only to aid visualization. Data from Bird 4.

Variation in wing shape is the geometric information that remains after accounting for stroke cycle kinematics within and among behaviour. In Figure 2, we visualise wing shape at selected stroke cycle phases, including during pronation, when the wing reverses from upstroke to downstroke; during the mid-downstroke; during supination, when the wing reverses from downstroke to upstroke; and finally, during the mid-upstroke. For example, rotating each wing configuration into the horizontal plane (without scaling, Figure 2 B i) reveals visual differences in the planform, some of which can be described by conventional shape descriptors. Other aspects of shape are more complex, as for example depicted by camber curves on the reconstructed wing surface at selected spanwise stations along the wing (Figure 2 B ii; wing twist removed to highlight camber variation at different phases of the stroke cycle).

**Figure 2.**
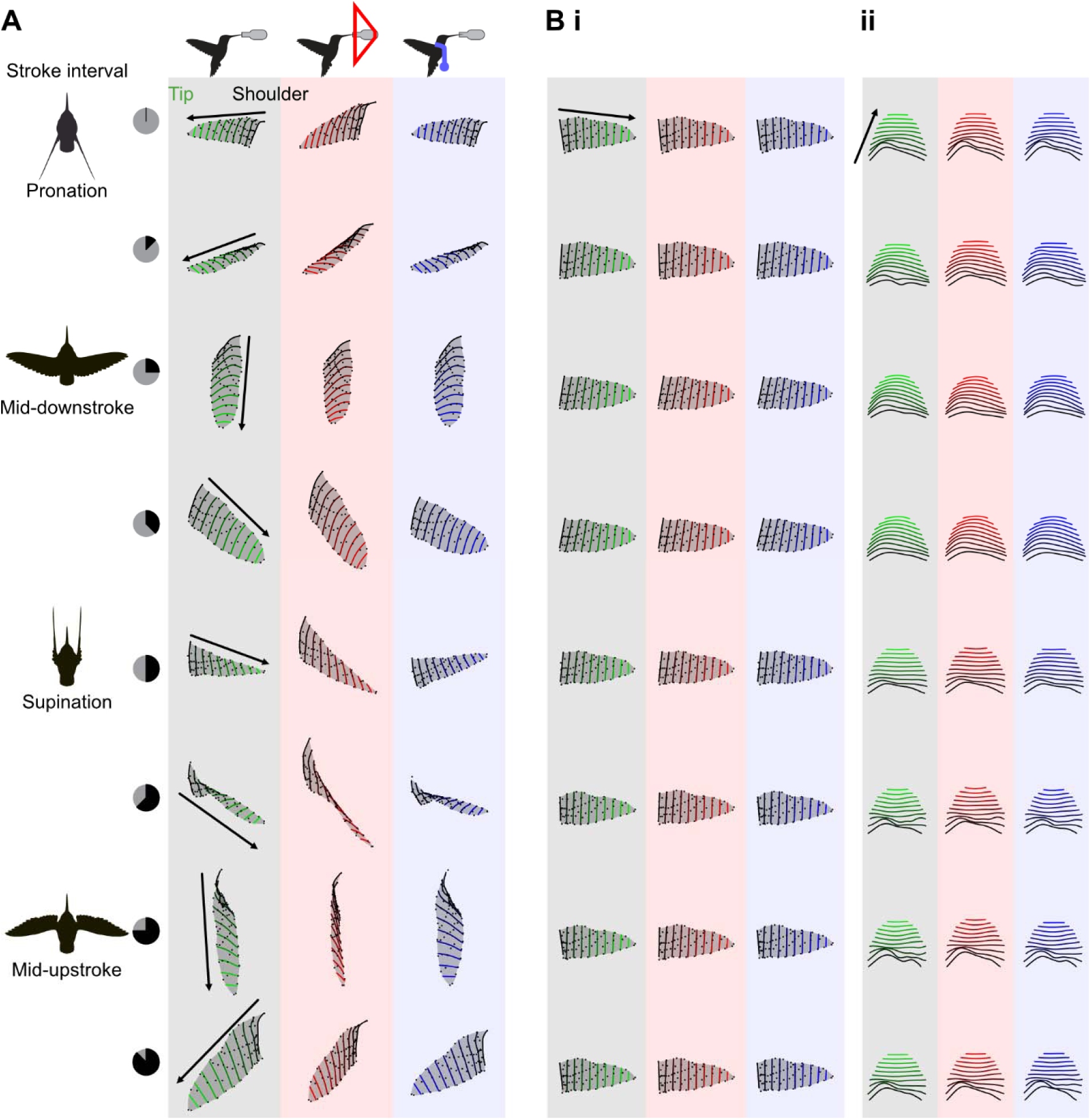
Exemplar hummingbird wing configurations throughout the stroke cycle. **A** Tracked landmarks (filled circles) were used to reconstruct the wing surface (chordal line segments) and its motions at eight selected time slices while hoverfeeding in typical conditions (black), in front of a visual mask (red), and during submaximal load lifting (blue). **B** Shape variation is revealed by eliminating rigid kinematic transformations between configurations to reveal differences in projected (*i*) and three-dimensional (*ii*) wing shapes. Wing twist is removed from *ii* to highlight camber variation and the sections are evenly spaced for visual clarity. Data from Bird 4.

Variation in kinematics and wing shape arises through concerted anatomical actions (Figure 3A). The anatomy suggests that the primary (P) flight feathers are actuated by either the carpometacarpus (P1– P6), or the first (P7–P9) or second (P10) phalanges of digit II. The secondary (S1–S6) flight feathers are actuated by the ulna. The contributions of these feathers to wing deformations throughout the stroke cycle are visualised through the approximately in- and out-of-plane trajectories of the tracked markers (respectively, *xy* and *xz* projections of the aligned configurations around this individual’s wing template, Figure 3 B). The *xy*-projected wing tip trajectories are nearly linear whereas those in the *xz* projection are circular, corresponding to the joint effects of wing twisting and wing folding and unfolding (area changes) throughout the stroke cycle (Maeda et al., 2017).

**Figure 3.**
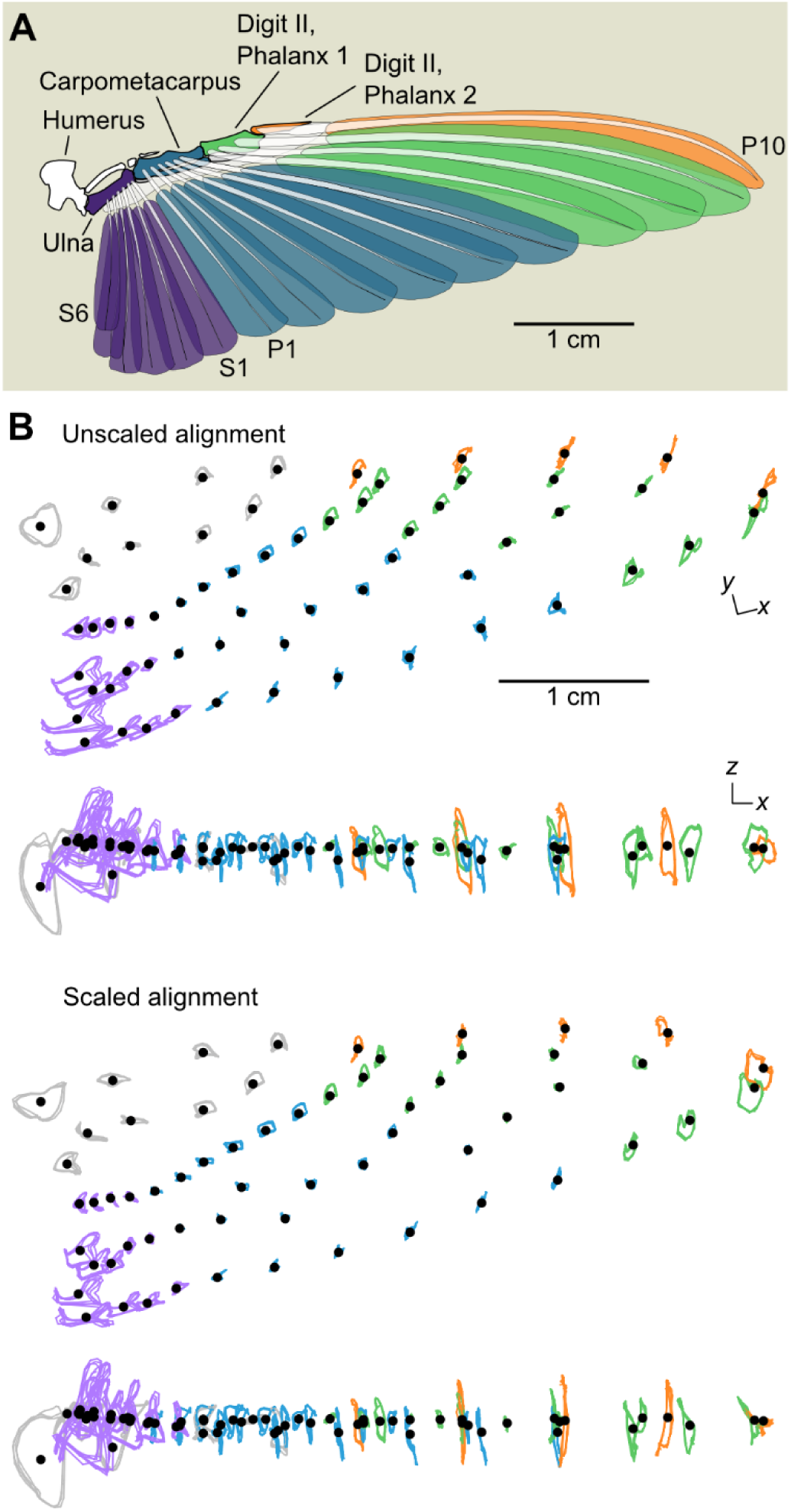
Functional anatomy of the hummingbird wing. **A** The hummingbird handwing is composed of four feather groups that insert on and are actuated by the ulna (secondary feathers, S6-S1), the fused carpometacarpus (primary feathers, P1-P6), and the first (P7-P9) and second (P10) phalanges of digit II. **B** Wing configurations at every time step were registered to the template configuration without scaling (top) or with scaling to unit centroid size (bottom). The trajectories of landmarks placed on the handwing (covert feathers) are depicted in gray, and others are coloured according to the scheme in A. Trajectories from Bird 4.

In our analyses, there are two important considerations of the alignment procedure. We align each configuration by all the landmarks. However, the wings are actuated by the handwing (Figure 3A), which does not lie at the landmarks’ centroid, and so centering induces unphysical apparent motions among aligned configurations (Figure 3). Aligning with all landmarks distributes tracking errors across the set.

Restricting alignment to the handwing markers (grey, Figure 3 B) may magnify spanwise errors because of the smaller alignment set that includes some markers with greater positional uncertainty, like the shoulder. This error will influence subsequent decompositions of the variance throughout the stroke cycle. A second consideration is the consequence of scaling configurations during alignment, which we visualise in Figure 3 B. Scaling transferred circular trajectories corresponding to wing folding to the *xy* projection (in-plane motions), whereas motions in the *xz* projection (out-of-plane) became more linear. In our view, scaling therefore preserves area changes as a source of approximately in-plane shape variation and allows us to attribute out-of-plane deformations to other causes as explored below.

### Morphological measurements

Morphological variation measured at equivalent stroke phases was generally low within individuals (mean mass and measurements in Supplementary Table 1). In two individuals, we also investigated the correspondence of in-flight measurements to those from posed wings with respect to the wing silhouette or the landmarks. Wing posing is typically intended to visually approximate in-flight wing shapes through feather spreading. Measurement of the landmark-defined wing perimeter in a manually posed wing yielded a similar wing area estimate in one individual compared to the mid-downstroke pose (−3% with respect to flight) but a large discrepancy in the other (−13%). The underestimated wing length in both individuals suggests this is a problem resulting from these birds’ very small wings and an obscured wing base while posing. A second and inherent problem is that wing surface area (and shapes based on it) will be underestimated when landmarks lie inside the wing perimeter rather than along it (compare posed measurements based on the silhouette or landmark perimeter, Supplementary Table 1). Nonetheless, the second moment of area 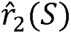, a measure of wing shape relevant to aerodynamics, was relatively insensitive to these errors in surface area and length measurement. Although this is not a detailed analysis, the sources of error suggest that discrepancies of ±15% between in-flight and posed measurements may be typical.

Hummingbird wings are highly dynamic during hoverfeeding (Figures 3 C and 4, Supplementary Table 2, S1 and S2 Videos). The surface area rapidly increases at the start of the stroke cycle (pronation) and reaches its maximum during the mid-downstroke (Figure 4 A). Due to wing expansion and folding, the second moment of area 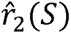 is greater (more of the wing area is distributed at a greater radius from the base) during the downstroke (wing expanded) than during the upstroke (partially folded; Figure 4 B). Wing twist, a function of the section angles of attack (Figure 4 F), was lowest during the downstroke (Altshuler et al., 2010a; Kruyt et al., 2014; Skandalis et al., 2017). The twist minima do not coincide with our placement of stroke reversals based on the wing azimuthal position. However, we note that some studies have demarked stroke reversals by the thinnest wing profile when viewed from a vertical position, corresponding to minimum twist (e.g., Altshuler et al., 2010a). The camber profile across the wing is complex (Figure 4 G), but greatest during the mid-downstroke, when the wing twists in the direction of feather reinforcement, resulting in a smoothly curved profile (see also Figure 2 B ii). During the upstroke, it appears that the ulna (based on Figure 3 A) cannot rotate sufficiently to maintain surface contiguity, which opens a gap between the primary and secondary feathers, as noted previously (Maeda et al., 2017). Our camber measurements most accurately reflect the true profile in the thin, distal wing sections, with some unaccounted-for ventral curvature due to flight feather P1’s thick rachis. At the wing base, the true camber profile is more complex because of the anatomical thickness profile (Figure 3 A). We therefore take the midspan camber profile as representative for subsequent analyses.

**Figure 4.**
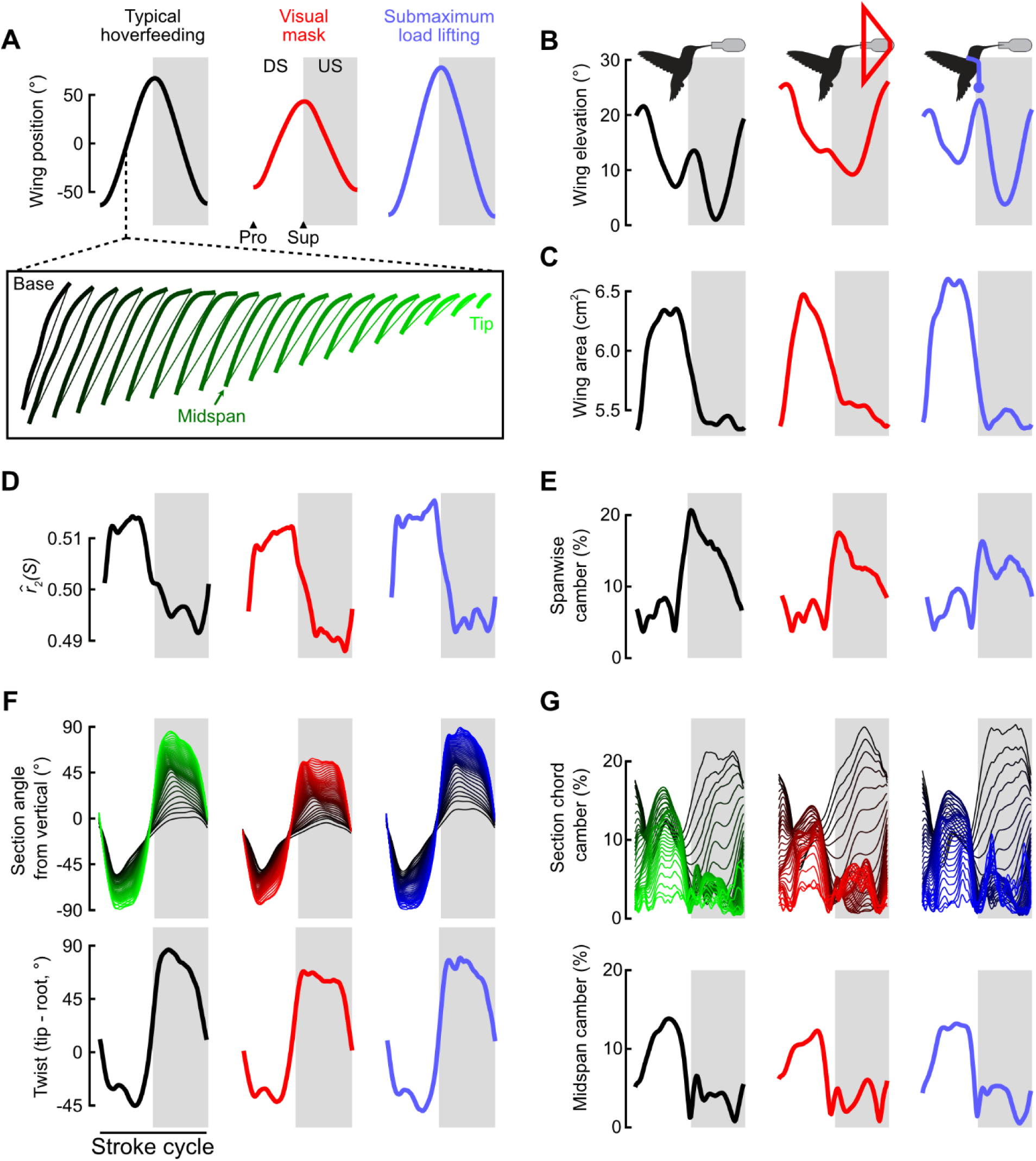
Kinematic and morphological variation among flight conditions in Bird 4. Each trace is the stroke-averaged response for one individual through downstroke (DS), supination (Sup), upstroke (US), and pronation (Pro). Box: Line segments and curves from a reconstructed surface used to quantify wing size and shape with spanwise segments colour-coded from black to green (typical hoverfeeding), red (visual mask), or blue (submaximum load lifting). Kinematic variation as quantified by the stroke amplitude (**A**) and elevation (**B**). Morphological variation as quantified by wing surface area (**C**), second moment of area 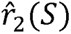, and spanwise camber (**E**). The sectional wing angle (**F**) and camber (**G**) profiles were used to derive the wing tip-to-base twist and midspan camber profiles, respectively.

Hummingbird wing size and shape were altered together with the kinematics when hoverfeeding in front of a visual mask or during submaximum load lifting (Mahalingam and Welch, 2013; Wells, 1993). In front of the visual mask, maximum wing area was increased during each of the mid-half strokes (downstroke and upstroke; Bird 4 presented in Figure 4 A). During submaximum load lifting, the wing area time series was similar to typical flight but with greater maximum area in both half-strokes. The second moment of area 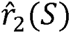 likewise differed between typical flight and either flight challenge in terms of the timing and amplitudes of identifiable peaks. Amongst the most notable differences were the twist and camber profiles in front of the visual mask. Wing twist during both half-strokes was substantially reduced, resulting from distal wing sections being much more inclined relative to the horizontal plane. In contrast, during submaximum load lifting the twist profile was similar to typical hoverfeeding but at a lower wing angle (Figure 2 A, 4 D). In some individuals, camber appeared to be slightly greater during both half-strokes whereas during submaximum load lifting the camber profile did not greatly differ.

Average measurements are reported in Supplementary Table 2. <u>Decomposition of dynamic wing shape</u> The statistical complexity of quantifying such complex spatiotemporal deformation patterns among behaviours through parametric descriptors alone motivated our approach to study variation through shape decompositions. Area changes across the whole wing, but so does the distribution of area with respect to the base. Many descriptors involve explicit choices, like reducing the wing sectional angle profile to a single measurement of twist or summarising the camber profile by its midspan measurement. Ostensibly, the spatial profiles of each could be highly relevant for describing variation within the stroke cycle and between behaviours. Some kinematic and morphological parameters at identified slices of the stroke cycle are summarised in Supplementary Table 2, but this approach could miss important modulatory intervals.

To compare among behaviours, we therefore used a data-driven approach to reduce the shape dimensionality and uncover shape variables that capture the major axes of variation in landmark configurations (Procrustes analysis). About 80% of the shape variance in the complete landmark set of each individual was recovered by including just 20 landmarks, whereas 95% recovery was achieved with 40-50 landmarks (Figure 5A). The asymptotic relationships suggest that, notwithstanding interindividual differences in marker number and placement, our marking consistently captured the range of shape morphological variation in typical hoverfeeding within each individual.

**Figure 5.**
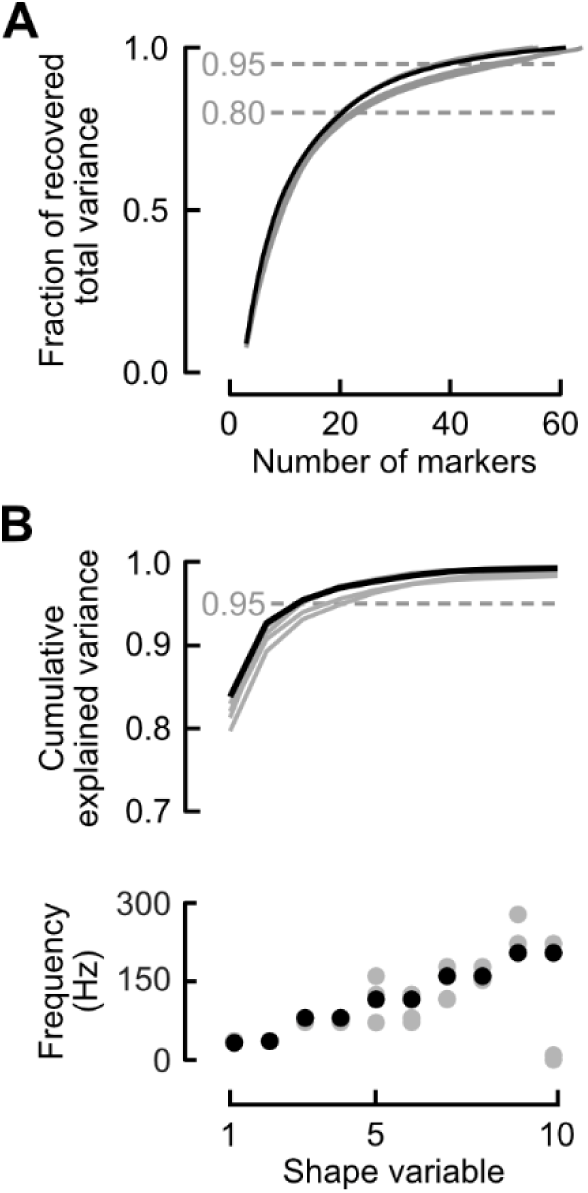
Linear decompositions of wing shape variation. **A** Minimal new shape information was gained beyond 40-50 markers in each individual. **B** The first four shape variables explained more than 95% of shape variance (top) and exhibited dominant frequencies at multiples of the stroke frequency (bottom). In all plots, each line or circle represents one individual, with Bird 4 highlighted.

The shape variables decompose periodic variation, and we therefore examined the dominant frequencies in each. We found that the shape variables, ordered by their singular values, were paired by their dominant temporal frequencies of integer multiples of the stroke frequency (frequency of the peak power/frequency in Figure 5 B; using unscaled alignments only slightly altered this result, analysis not shown). This was most evident in two individuals, in which at least the first 10 shape variables occurred in frequency pairs; the addition of two more cameras for these individuals may have improved our power to identify the dominant frequency in low-variance shape variables. Nonetheless, all individuals consistently exhibited dominant frequency pairs in the first four shape variables, and we therefore focus on these four, which respectively explained ∼82%, 10%, 3%, and 1.5% of total shape variation (Figure 5 B). The spatiotemporal structure of deformations represented by the shape variables was studied through reduced-rank reconstructions of the wings based on the extremal shape scores. In Figures 6 A-D i, an exemplar template geometry is plotted together with vectors representing each landmark’s displacements along each shape variable’s axis (Bird 4; Figure 6 A-D i, vector length doubled for visualization; Figure 6 A-D ii, shape scores). Visually, these vectors suggest out-of-plane (first and third shape variables) and in-plane (second and fourth) deformations, supported by the average angle of the landmark vectors with respect to the horizontal plane (Figure 6 A-D iii).

**Figure 6.**
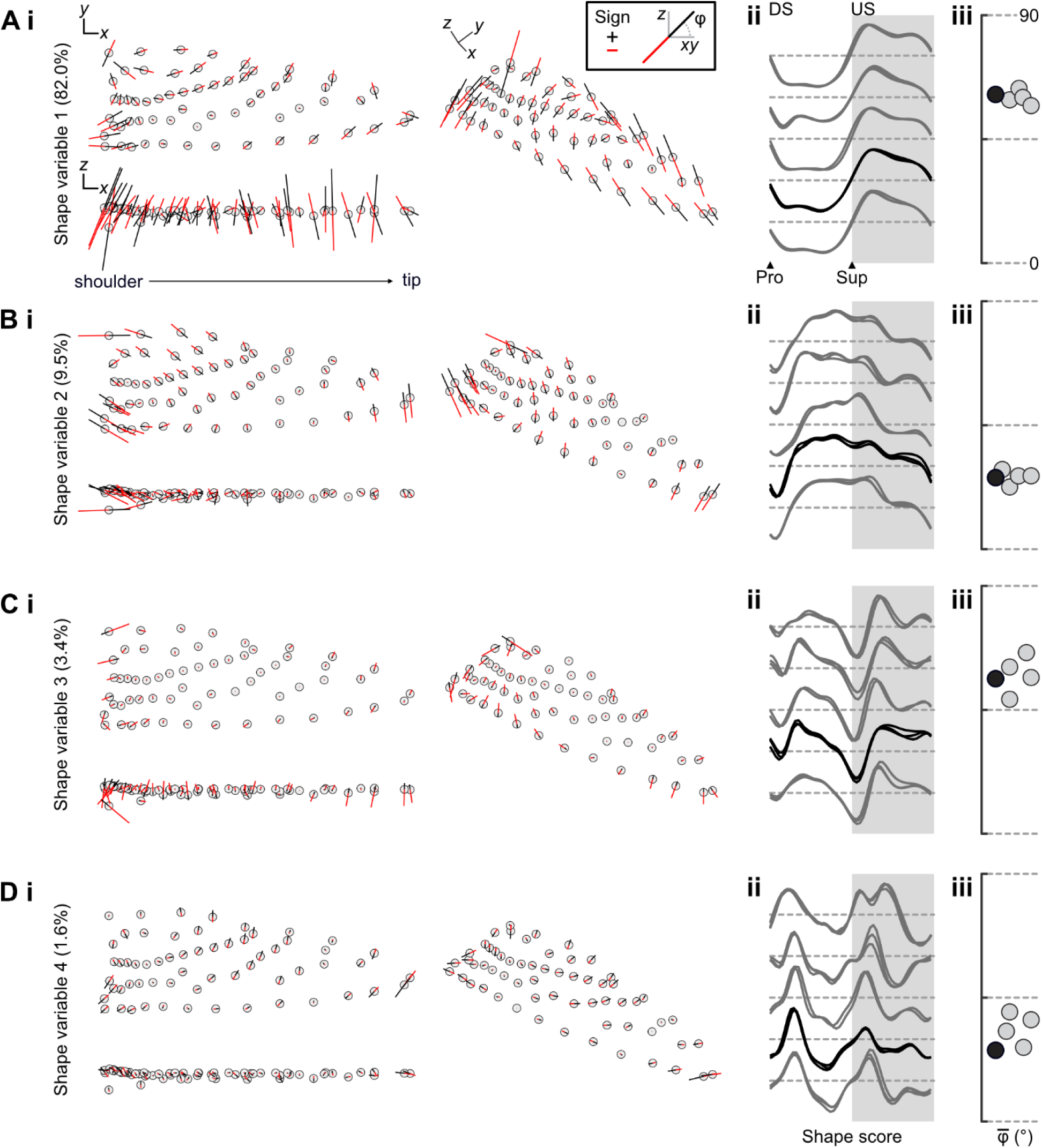
Shape variables quantify in- and out-of-plane deformation components. **A-D** *i* Marker (open circle) trajectories along each shape variable’s axis, plotted as vectors spanning minimum (black) to maximum (red) shape scores (scaled to twice this individual’s range of variation). *ii* Normalised stroke cycle shape scores per individual. *iii* Average vector angle <-of trajectories with respect to the horizontal plane. Representative configurations and individual traces from Bird 4 in black, others grey.

The temporal evolution of shape scores was highly consistent across individuals in each shape variable. The spatial pattern of out-of-plane deformations represented by the first shape variable points to wing twisting in phase with the mid-strokes (mid-downstroke and mid-upstroke). Conversely, scores on the second shape variable, representing primarily in-plane deformations, rapidly increased from negative to positive through the downstroke. The scores stayed elevated during supination and the early upstroke, after which they fell again to negative values. This suggests the second shape variable particularly captures the folding and unfolding of the wing during reversals, mostly out of phase with the first shape variable. The third and fourth shape variables complemented the deformations represented in the first two, at twice the frequency. The third shape variable contrasted out-of-plane deformations that were most distinct during supination and the midstroke poses. The fourth shape variable contrasted in-plane deformations that were most distinct in the early and late downstroke, and in the early upstroke. Scores in the upstroke were less consistent among individuals. This could be because shape variables explaining smaller amounts of variance may be more influenced by individual deformation variation or by the marking schemes, which we cannot distinguish here. Altogether, we interpret each shape variable frequency pair, ranked by explained variance, as contrasts between the sources of in- and out-of-plane at identifiable intervals of the stroke cycle.

### Interpreting shape variables through wing descriptors

We next sought to understand the correspondence of our parametric descriptors (Figure 4) to the major variation captured by shape variables, and equally to describe the shape variables in real terms. We first examined the cross-correlation of each shape variable with key kinematic and morphological parameters. The first shape variable was a quarter cycle out of phase with the azimuthal wing position, whereas the second shape variable was in phase (Figure 7). These phase relationships suggest the first and second shape variables respectively encompass shape variation occurring at stroke midpoints and reversals. The third and fourth shape variables were better associated with wing elevation, which is similarly twice the stroke frequency for a typical J-shaped wing tip path (Figures 1, 4 B). Among morphological descriptors, wing twist was the sole parameter to be strongly linearly correlated with a shape variable (Figure 7). The other variables exhibited complex periodicity. Taking the lag into account through cross-correlation analysis, wing area and chordwise camber were the best correlated with the second shape variable. Spanwise camber was the sole descriptor to exhibit nearly in-phase relationships with the third shape variable. There were no obvious relationships between the fourth shape parameter and any shape descriptor when examining each relationship alone.

**Figure 7.**
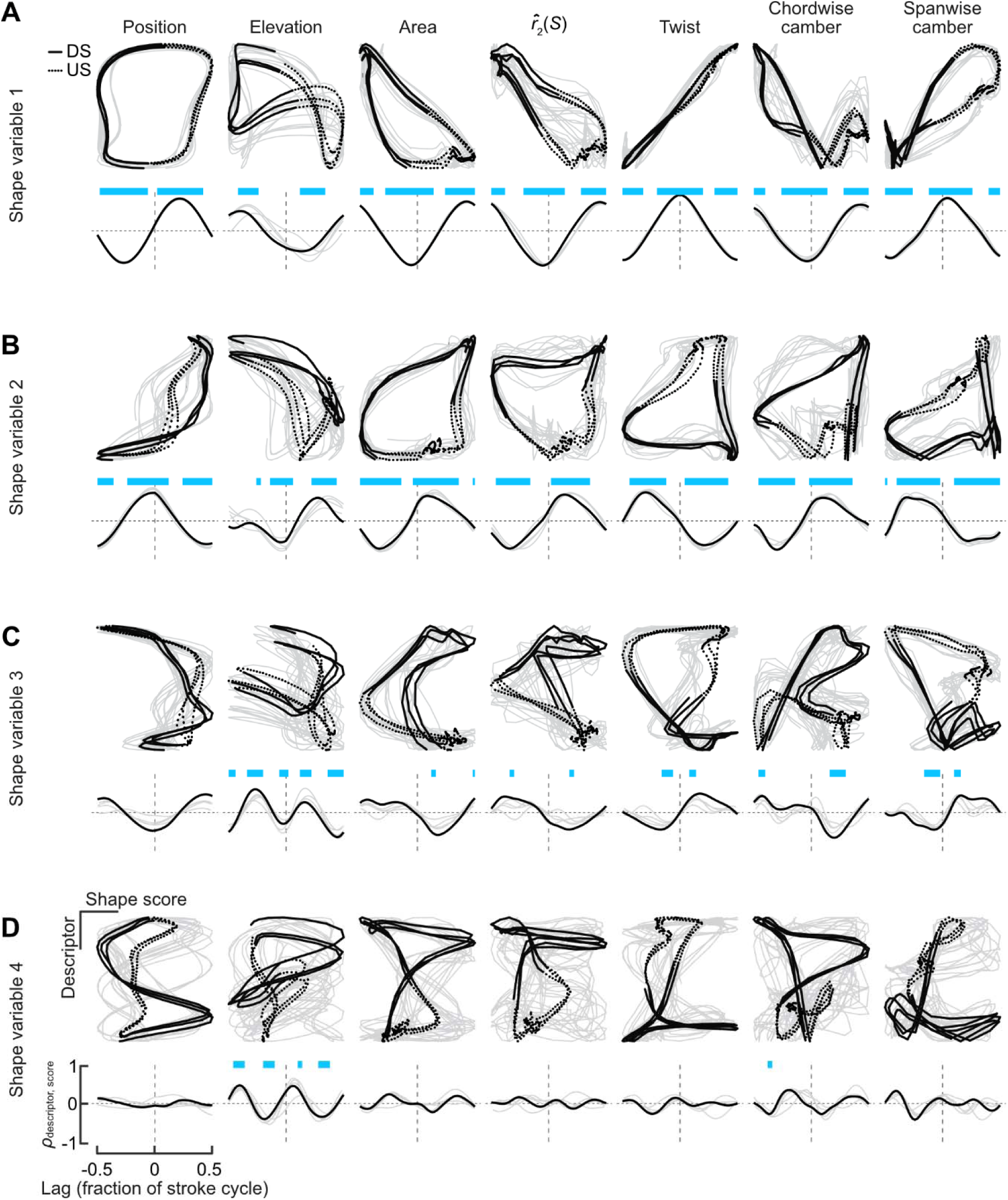
Cross-correlations between shape variables and parametric descriptors of the wing configuration. **A-D** *Above* Plot of each relationship through multiple stroke cycles (DS: downstroke, solid line; US: upstroke, dashed line). *Below* Cross-correlation coefficient ρ_descriptor,score_ at each time lag as a fraction of the stroke cycle. Lags of 0.25 and 0.50 corresponds to quarter- and half-cycle differences. Time lags with significant correlations are highlighted in blue. Representative data from Bird 4 in black, others grey.

We applied multiple regression models to identify relationships among shape scores and wing descriptors. We included kinematic parameters to account for mutual associations with an underlying periodic signal rather than any direct relationship. The first shape variable was almost entirely explained by variation in wing twist (normalised regression weights β in Figure 8), reinforcing our visual interpretations based on the marker motions (Figure 6) and cross-correlation analysis. Moreover, the model *R*^2^ (shape descriptors together with kinematics) of the first shape variable was close to unity.

**Figure 8.**
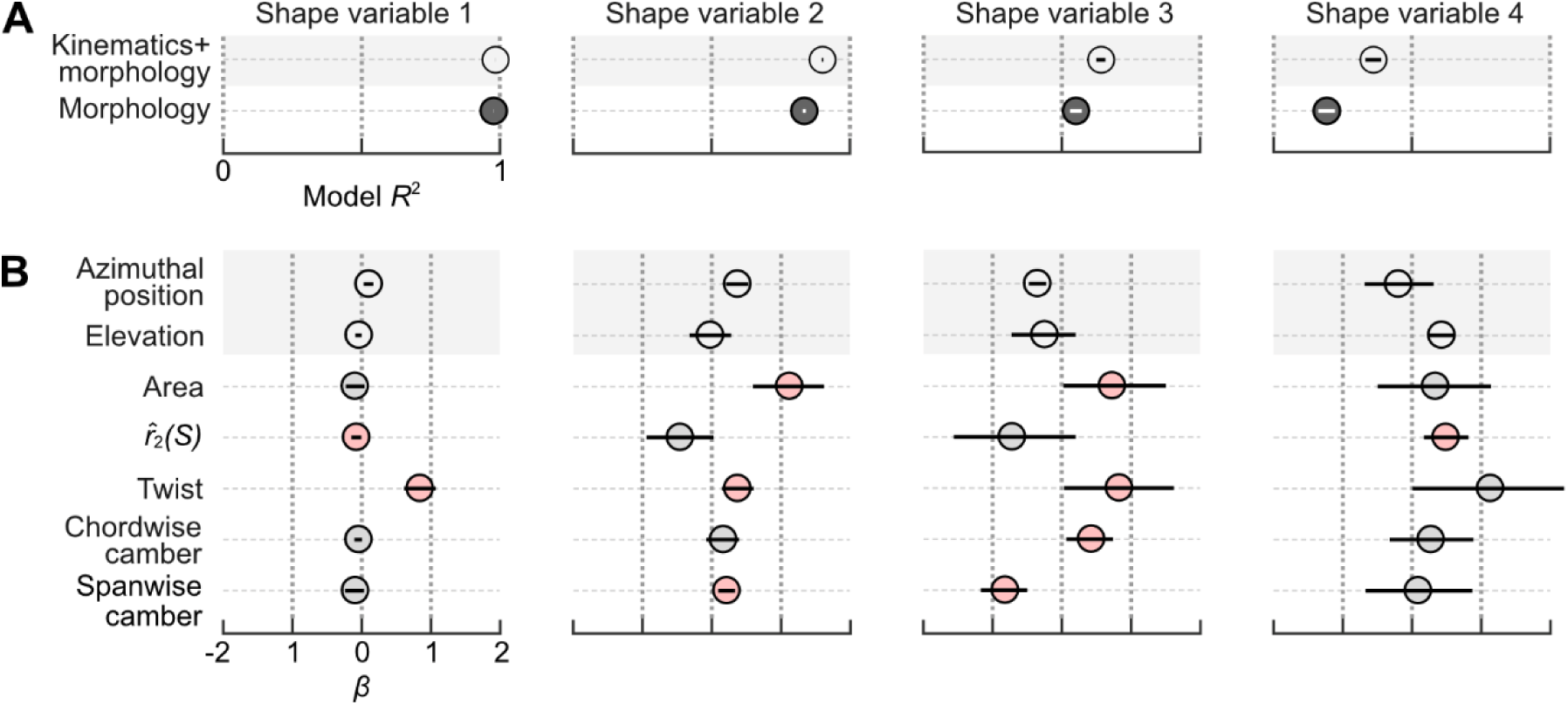
Multiple regression coefficients of shape variables with parametric descriptors of wing configurations. **A** Model *R*^2^ including or excluding kinematics. **B** Standardised regression coefficients β for each model parameter. Credible intervals of parameters in red exclude zero. Second moment of area, 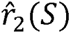.

Unlike the first shape variable, none of the other shape variables could be considered simple linear functions of morphological parameters, because none were singly correlated with near-zero phase (Figure 6). The second shape variable, representing in-plane deformations, was best associated with temporal trends in area, even though the dynamics of the respective variables differed, especially during reversals (compare to Figure 4). This relationship likely arises from our scaling of the configurations during alignment, as observed by inspecting the impact on the aligned marker trajectories (Figure 3 B). The third and fourth shape variables were less well explained, with model *R*^2^ ∼ 0.64 and 0.36, respectively. All morphological parameters other than 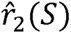 were associated with variation in the third shape variable, although excepting spanwise camber, each had wide credible intervals spanning near zero. Based on these linear associations and the predominance of out-of-plane deformations along the span, we therefore attribute the third shape variable primarily to spanwise deformations. The sole parameter associated with the fourth shape variable was 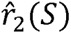. As the fourth shape variable likewise relates to in-plane deformations, we describe it as representing shape variation related to the distribution of wing area.

### Spatiotemporal variation in wing shape among hoverfeeding behaviours

Pre-selecting time slices at which to describe variation in wing size and shapes, such as during the mid-strokes or reversals (Supplementary Table 2), may reduce our power to detect real differences in flight behaviour. We applied statistical splines (Wood, 2017) to identify temporal intervals in which the shape variable scores significantly differed between typical hoverfeeding and the two challenged flight conditions. Changes in shape score were identified with respect to the mean time series of typical hoverfeeding (Figure 9), either during specific temporal intervals or throughout the stroke cycle (Figure 9 B). These differences are summed to illustrate the overall changes in shape scores in each flight condition (Figure 9 C). Individual random temporal variation was evenly distributed around zero over the course of the stroke cycle (Figure 9 D), suggesting there was no systematic underfitting of the population model at any point.

**Figure 9.**
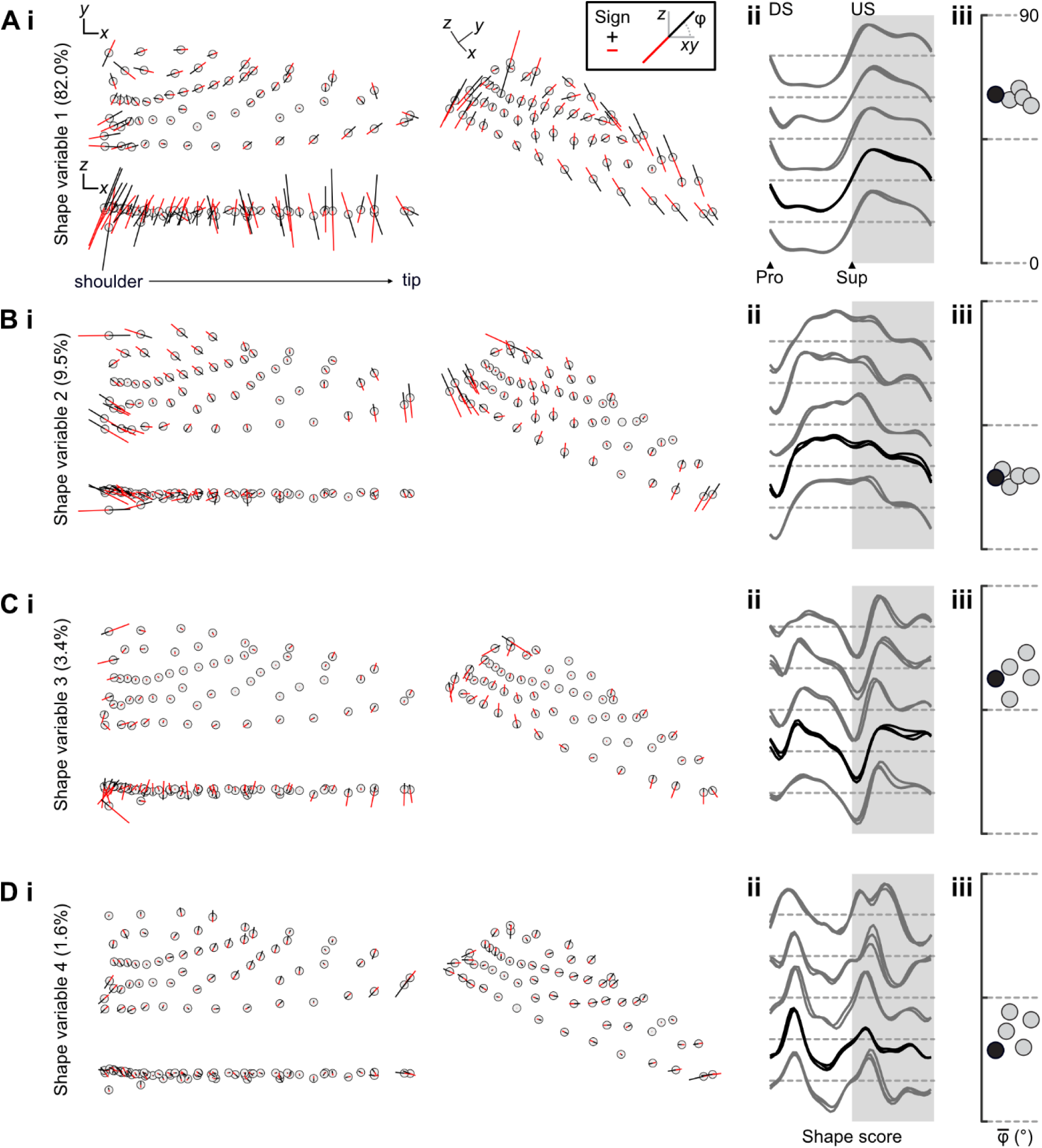
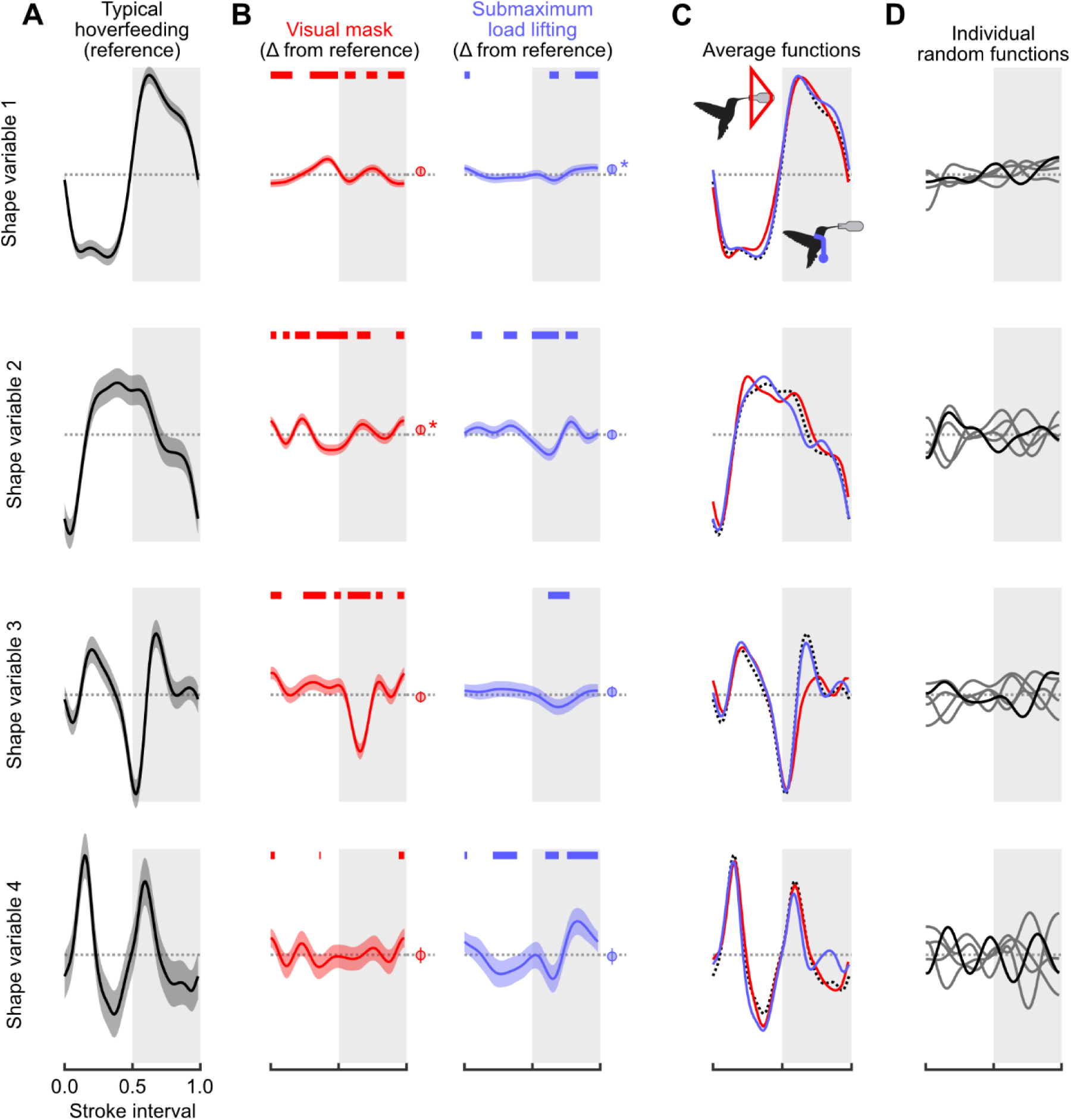
Hoverfeeding in different flight conditions is associated with distinct wing spatiotemporal deformation dynamics. **A** Reference time series of shape scores in typical hoverfeeding, representing typical wing deformations. **B** Time-dependent and global differences in the time series of shape scores in challenged flight with respect to typical hoverfeeding (Δ from reference). Thick overlines denote temporal intervals that significantly differ from the reference. Average shape score differences (intercept difference) are shown to the right with an asterisk where the credible intervals exclude zero. **C** Summing the reference function (A) with the non-linear difference functions and parametric intercept (B) shows the mean shape trends in each treatment condition. **D** Random temporal effects for each individual do not exhibit major trends that would suggest modelling biases. Scale is constant in all panels.

The time series of shape scores in each challenged flight condition point to distinct spatiotemporal deformation patterns. The nonlinear component of the time series of shape scores in front of the visual mask suggests a contribution of wing deformations throughout the stroke cycle. Scores of the first shape variable suggest a reduced wing twist (shape score closer to zero) during the late downstroke and mid-upstroke, and greater twist during the mid-upstroke. Prominent differences in the time series of the second shape variable scores point to widespread changes in the wing area dynamics particularly coinciding with the midpoints of the down- and up-strokes and the stroke reversals. A slight intercept shift on the second variable with respect to typical flight also suggests that wing area may be slightly greater overall in front of the visual mask. The time series of the third shape variable scores exhibited the most dramatic changes in shape scores relative to typical flight, as a large dip during the upstroke effectively offset the peak during the same phase of typical hoverfeeding (visible in Figure 9 C). There were some identifiable differences of the time series of the fourth shape variable during pronation, but these were of small effect. During submaximum load lifting, the time series of shape scores instead points to a prominent role of in-plane deformations, particularly during the latter downstroke and throughout the upstroke. Scores on the first and third shape variables did not greatly differ from typical hoverfeeding except for the early upstroke and a large interval encompassing the late upstroke and pronation.

Conversely, the second and fourth shape variables suggest widespread and prominent changes in area dynamics during the late downstroke and throughout the upstroke, coinciding with observed changes in wing area and 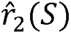 (Figure 4).

An important observation from these analyses is that the dynamic shape variation among behaviours is not exclusively contained on the first shape variable, even though it explains the majority of shape variation in typical hoverfeeding. The third and fourth shape variables appear to identify modality-specific deformation patterns that are not very well explained by parametric descriptors and which each explain <10% of shape variation. We thus argue that explained variance (ranking of the squared singular values) is insufficient for predicting the sources of behavioural plasticity. Important behavioural modulation may be captured in low-variance components, corresponding to subtle types of spatiotemporal deformations.

## DISCUSSION

Behavioural flexibility in body shape blurs the distinction between kinematics and morphology. Body deformations give rise to fluid-structure interactions that directly influence the forces produced during locomotion (Boerma et al., 2019; Cheney et al., 2020; Ravi et al., 2015; Ravi et al., 2020; Standen and Lauder, 2005; Swartz et al., 2006). The contributions of wing deformations to hummingbirds’ agile flight has been unclear (Kruyt et al., 2014; Skandalis et al., 2017; Tobalske et al., 2007). Some of the complexity arises from tracking and comparing shape dynamics among flight behaviours in what has generally been perceived as a stiff, insect-like wing with primarily kinematic force modulation. We developed a weightless and minimally disruptive wing marking procedure (Figures 1-3) to capture the spatiotemporal dynamics of wing deformations in Anna’s hummingbird (Figure 4). We identified major axes of shape changes (Figures 5, 6) and their relationships to conventional descriptors (Figures 7, 8). On this basis, we investigated the timing and origins of differences in wing shape between typical and kinematically constrained hoverfeeding (Figures 1, 9). Distinct sources and timings of deformations were associated with each. We consider our findings in the context of the level of detail required for analyses of functional morphology.

How many shape variables are needed to explain hummingbird wing shape variation? In similar analyses with other animals, the number of preserved components has been chosen statistically or as a threshold of the explained variance, but neither method relates directly to the shape variation (Bartol et al., 2018; Bozkurttas et al., 2009; Dryden and Mardia, 2016; Riskin et al., 2008; Schlager, 2017). Our results lead us to deemphasise explained shape variance as a measure of shape variable importance. In our study, the first two shape variables represented ∼90% of the shape variation. However, we found that the shape variables were ordered by their dominant frequencies, which offered a principled method for selecting the number to preserve for further analysis. We studied the first two integer multiples of the stroke frequency, comprising in- and out-of-plane deformation components (higher multiples had higher uncertainty in some individuals, Figure 5). Combining parametric and non-parametric approaches allowed us to interpret these deformation modes in real terms. Altogether, this method for retaining four shape variables allowed us to study changes in wing shape through temporal contrasts of in- and out-of-phase deformations at two frequencies.

The first two shape variables explained the majority of variation throughout typical stroke cycles and corresponded to important descriptors of wing size and shape. Hummingbirds’ chained joint rotations enable wing supination, upstroke weight support, and sustained hovering [Figure 3, (Hedrick et al., 2011; Warrick et al., 2005)]. Accordingly, we found that the majority of wing shape variation (∼80%) throughout the stroke cycle was visually and linearly associated nearly exclusively with wing twisting. The converse was not true, and twisting was associated with other shape variables as well (Figure 8). One explanation is that our twist measurement correlates broadly with out-of-plane deformations, especially the related or induced deformations like chordwise cambering and feather separation (Ennos, 1988; Maeda et al., 2017). If so, this suggests that our approach allows us to identify the influence of wing twisting (the major actuation mode) on the various induced deformation modes. The second shape variable was primarily related to the wing area dynamics. This specific ranking followed from the scaling of each configuration during the alignment step, which we found preserved the sense of area changes as a component of in-plane deformations (Figure 3 B). Otherwise, area would correlate with the first shape variable, as expected (analysis not shown; Dryden and Mardia, 2016; Schlager, 2017). These area changes corresponded to ∼10% of the stroke cycle shape variation and point to a more limited capacity to modulate the wing’s dimensions than its rotation (Hedrick et al., 2011; Tobalske et al., 2007). The second two shape variables explained deformations at a higher frequency than the first two, which explains their smaller partitionings of shape variance. Higher frequency components naturally correspond to shorter intervals with fewer sampled configurations. For example, contrasts between the start and end phases of the downstroke are particularly evident in the fourth shape variable. Although these shape variables represent higher frequency deformations, we still found they correspond to functionally relevant descriptors of wing shape. We attribute the third shape variable to this species’ limited potential for spanwise deformations, together with correlated changes. Despite an overall low model *R*^2^, spanwise camber was prominent in the prediction of third shape variable scores (Figure 8). Our interpretation of a spanwise mode was supported by visual inspection (Figure 6) and cross-correlation (Figure 7). Alternative parametrisations of spanwise bending might capture other spanwise information, possibly with higher correlation and different phase. If so, this might suggest a method for identifying sets of strong parametric descriptors based on their relationships to the shape variables. We attributed the fourth shape variable to in-plane shape variation with a weak association to the second moment of area 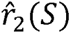 (Figure 8). Similarly to the arguments for spanwise bending, it may be that the fourth shape variable is related to an ‘area distribution mode’ that is only weakly identified by 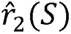. This weak identification is somewhat supported by the observation that the second moment of area is overall fairly insensitive to many changes in shape (Figure 4, Supplementary Table 1).

The shape variables’ frequency and phase relationships allowed us to identify spatiotemporal differences in wing deformations that would not be predicted simply by the fraction of explained variance alone. Put another way, the axes that explain the most variance within typical hoverfeeding behaviour do not necessarily predict the sources of variation among behaviours. Despite explaining the majority of shape variation, differences in shape scores on the first shape variable between typical hoverfeeding and in front of the visual mask were of generally small magnitude throughout the stroke cycle. During submaximum load lifting, differences were restricted to the upstroke and pronation. Conversely, relative to the scale of shape scores in typical hoverfeeding (Figure 9 A), more substantial changes were represented on each of the other shape variables despite accounting for 10% or less of typical hoverfeeding shape variance. We cannot discount that noise, including individual variation, marker errors, and temporal alignment, becomes increasingly influential in shape variables explaining less variance. An alternative explanation is that because the first shape variable explains the most variance, it may also be the most functionally constrained. Wing twist is necessary to create a lift-generating conformation with an appropriate angle of attack in both the down- and upstrokes (Altshuler et al., 2010b; Hedrick et al., 2011; Song et al., 2014). As the majority of force production depends less on the details of other aspects of size and shape (Figure 5), like wing area and bending, these may therefore have a greater scope for force modulation if actively controlled. Similarly, the first shape variable is in phase with the wing sweep (Figure 6), whereas the other shape variables explain less overall variation but encompass important intervals like stroke reversals. Shape variation in these intervals may have an outsized effect on aerodynamic forces (Dickinson et al., 1999; Haque et al., 2023; Ravi et al., 2020; Song et al., 2015). The relative significance of changes in each shape variable can be studied through computational and robotic simulations (Akanyeti et al., 2016; Bozkurttas et al., 2009; Zheng et al., 2013).

Alternative methods for investigating the variation in the shape space could offer other insights and reveal other details. Factor analysis may reveal latent factors corresponding to particular feather groups (Figure 3) or parametric wing descriptors. Dynamic mode decomposition could be used to factorise the linear and temporal sources of shape variation (Brunton et al., 2016). What shape changes are revealed crucially depends on the shape metric. The Procrustes metric applied here is well explored in the analysis of static shapes (Dryden and Mardia, 2016; Rohlf, 2000; Slice, 2001) but has received less attention in time series of shapes and particularly of periodic motions. Some effects of decisions of this alignment procedure were highlighted in Figure 3. We anticipate theoretical developments in computational geometry and metrics for comparing time series of flexible bodies.

Detailed analyses of flexible bodies have provided important insights into the evolution of morphological structures and animal performance (Lehmann et al., 2011; Lucas et al., 2014; Mountcastle and Combes, 2013; Young et al., 2009; Zheng et al., 2013). In the present study, the sufficiency of the time-consuming measurements could only be examined *post hoc* (Figure 5). Detailed surface reconstructions remove some elements of choice in terms of feature tracking and labour costs, but shape analysis through point clouds and surfaces is considerably more complex (Beg et al., 2005; Qiu et al., 2009). We offer some viewpoints when considering the detail with which wing size and shape is quantified for the purposes of ecological and evolutionary inference.

Wing posing is intended to broadly emulate wing shapes in flight (Supplementary Table 1). We found a strong correspondence between the dynamics of the biomechanically motivated parametric descriptors and the non-parametric approaches to measuring shape (Figure 8). This suggests that such descriptors will continue to have a strong role in comparative inference, including on static specimens. It is increasingly feasible to measure functionally critical three-dimensional shapes, like wing camber, in static specimens (Rader and Hedrick, 2023). As we observed here, wings assume a roughly similar size and shape during much of the stroke cycle and between successive cycles (Supplementary Table 1).

Accordingly, the scores of the first shape variable quickly stabilised and were maintained for most of the downstroke. The central question is the extent to which the posed measurements reflect real in-flight postures. Although wing shapes substantially depend on bone and feather geometry, nearly all *measurements* of shape depend on posing. Feather spreading by posing (Supplementary Table 1) or musculoskeletal actuation should be carefully considered, and care should be taken that the posing itself does not induce the shape variation (bias). Our measurements suggest that errors up to 10-15% may arise through posing (Supplementary Table 1). However, the hummingbirds’ small size was certainly a factor and so this estimate may be an upper limit to that in larger birds. Where possible, we encourage repeated measurements, including among experimenters to capture different conceptions of flight-like poses. Intraspecific variation, including measurement error, can be influential in comparative analyses [see refs in (Skandalis et al., 2017)], and errors may differ among measures of size and shape.

Although static measurements remain an important resource, evolution acts on body dynamics that are not quantified by posing. It remains unclear to what extent variation in morphology predicts aspects of behavioural variation like agility rather than the demands of mean weight support (Baliga et al., 2019; Dakin et al., 2018). The significance of shape variation that arises through environmental interactions, like ubiquitous tip bending (Lucas et al., 2014), must be measured in behaving animals. The most detailed analyses will be expected of animals with soft bodies (Cheney et al., 2022; Gemmell et al., 2015; Lehmann et al., 2011; Muijres et al., 2008) and complex force generation patterns (Bartol et al., 2018; Dickinson et al., 1999; Ravi et al., 2020; Warrick et al., 2005). However, as the degree of body stiffness is continuous, we believe an important tool can be quantifying the prevalence of deformations through their frequency spectrum. Relatively rigid bodies must have little or no power at any frequency since differences in configuration can be explained through rigid transformations rather than shape changes. A broader frequency spectrum signifies a wider range of spatiotemporal deformation scales related to locomotor cycles (Figures 5, 6) and to effects like aeroelastic flutter and vortex induced vibrations. By compressing shape changes into fewer variables, it becomes possible to apply familiar tools to analyse and compare time series in order to investigate variation among individuals and species.

Shape spaces provide a natural setting for studying the control and evolution of biological deformations [see, for example, the review of closed-loop control by (Roth et al., 2014)]. To this end, we require detailed knowledge of how configurations and changes in configurations depend on neuromuscular signals and environmental feedback. This necessarily entails pairing deformation studies with the actuating movements (Figure 3; Hedrick et al., 2011) to parse the muscular, inertial, and environmental forces acting on the wing (Badger et al., 2019; Haque et al., 2023; Song et al., 2014). In this respect, the shape components may prove to be particularly useful for delineating notions of coarse and fine motor control in continuously deforming bodies, as may be represented through our shape variables. In hummingbirds, coarse control through power muscle activity (pectoralis major and supracoracoideus) has been well-studied (e.g., Altshuler et al., 2010b; Konow et al., 2017; Mahalingam and Welch, 2013; Ravi et al., 2020), but the roles of handwing muscles in fine control remain largely enigmatic (Altshuler et al., 2012). Fine muscular control may not only generate deformations but also resist them through stiffening that limits the influence of aerodynamic and inertial forces, as has been observed in fish bodies (Tytell et al., 2016). In our experiments, handwing muscles might have been recruited to act against inertial bending due to the rapid accelerations of the foreshortened stroke cycle in front of the visual mask. The handwing muscles could also be involved in hummingbird inertial steering maneuvers more broadly (Haque et al., 2023). Understanding the fluid-structure interactions and how specific deformations interact with fluid flows and kinematics will be helped by detailed body and wing surface reconstructions (Cheney et al., 2020) together with computational methods for registering such detailed geometry (Beg et al., 2005; Qiu et al., 2009). Altogether, we expect that detailed shape analysis and shape spaces will have an important role in uncovering functional and evolutionary principles of soft bodies and their complex dynamics.

## ACKNOWLEDGMENTS

We are especially grateful to Dr. Laurent Younes for insightful discussions on quantifying dynamical shape variation, and to Drs. Rajat Mittal and Daniel K. Riskin on flapping aerodynamics.

## DATA AVAILABILITY

The data have been deposited in the figshare repository at doi:10.6084/m9.figshare.23234915. Software code is available upon reasonable request.

