## Supplementary Materials for "The spatiotemporal richness of hummingbird wing deformations"

**Supplementary Table 1** Average morphological measurements of the five individual hummingbirds in this study. Values in flight are presented as means and 95% confidence level. For individuals 4 and 5, we also report measurements from posed specimens based on the wing’s silhouette or the landmarks on the wing perimeter, together with the percentage difference from the in-flight measurement. Non-dimensional second moment of area, *r̂*_2_(*S*).

| **Bird** | **Body mass (g)** | **Length (cm)** | **Area (cm^2^)** | ***r̂*_2_(*S*)** |
| --- | --- | --- | --- | --- |
| 1 | 4.08 (0.03) | 4.969 (0.008) | 6.60 (0.02) | 0.503 (0.003) |
| 2 | 3.78 (0.01) | 5.128 (0.002) | 6.60 (0.01) | 0.497 (0.001) |
| 3 | 3.93 (0.04) | 4.909 (0.006) | 6.05 (0.02) | 0.492 (0.001) |
| 4 | 4.11 (0.04) | 4.972 (0.004) | 6.34 (0.03) | 0.501 (0.001) |
| 4 (posed silhouette) |  | 4.855 (-2.4%) | 6.85 (+10%) | 0.491 (-2.1%) |
| 4 (posed markers) |  | 4.733 (-4.8%) | 6.12 (-3.5%) | 0.483 (-3.6%) |
| 5 | 4.18 (0.02) | 5.027 (0.007) | 6.35 (0.03) | 0.487 (0.001) |
| 5 (posed silhouette) |  | 4.633 (-7.8%) | 6.17 (-2.8%) | 0.498 (+2.2%) |
| 5 (posed markers) |  | 4.552 (-9.5%) | 5.53 (-13.0%) | 0.481 (-1.3%) |

**Supplementary Table 2** Kinematic and morphological variation among hummingbirds during identified phases of three flight conditions. Values are presented as means and 95% confidence level for the number of individuals (N) in each condition. Midspan section angle relative to vertical, α. Non-dimensional second moment of area, *r̂*_2_(*S*).

| **Stroke phase** | **Condition** | **N** | **Position (°)** | **Elevation (°)** | **α  (°)** | **Length (cm)** | **Area (cm^2^)** | ***r̂*_2_(*S*)** | **Twist (°)** | **Chordwise camber (%)** | **Spanwise camber (%)** |
| --- | --- | --- | --- | --- | --- | --- | --- | --- | --- | --- | --- |
| **Pronation** | Visual mask | 4 | 49.12  (5.697) | 19.347  (3.854) | 76.422  (2.846) | 4.722  (0.04) | 5.455  (0.147) | 0.487  (0.004) | 4.58  (3.597) | 5.248  (0.718) | 6.455  (0.637) |
|  | Typical flight | 5 | 28.388  (7.415) | 22.239  (2.392) | 83.548  (1.917) | 4.749  (0.048) | 5.47  (0.12) | 0.485  (0.003) | 7.472  (2.781) | 4.321  (0.731) | 6.954  (0.719) |
|  | Submaximum load lifting | 3 | 24.881  (10.332) | 23.123  (2.031) | 84.26  (1.834) | 4.752  (0.012) | 5.452  (0.069) | 0.485  (0.003) | 6.485  (3.534) | 5.18  (0.62) | 7.123  (1.096) |
| **Mid-downstroke** | Visual mask |  | 93.185  (5.5) | 9.439  (5.566) | 29.593  (3.014) | 4.96  (0.062) | 6.414  (0.188) | 0.499  (0.003) | -36.98  (2.656) | 10.048  (0.447) | 7.391  (0.697) |
|  | Typical flight |  | 91.276  (5.475) | 7.392  (2.318) | 20.719  (2.275) | 5.001  (0.039) | 6.388  (0.106) | 0.496  (0.003) | -35.456  (1.665) | 12.033  (0.43) | 6.033  (0.565) |
|  | Submaximum load lifting |  | 98.195  (8.605) | 12.79  (1.261) | 19.866  (3.813) | 5.031  (0.015) | 6.5  (0.054) | 0.497  (0.004) | -34.942  (3.907) | 12.537  (0.253) | 7.837  (0.624) |
| **Supination** | Visual mask |  | 131.825  (4.284) | 2.994  (6.401) | 74.789  (2.809) | 4.79  (0.038) | 5.75  (0.118) | 0.486  (0.005) | 59.1  (6.204) | 5.143  (1.318) | 19.9  (0.918) |
|  | Typical flight |  | 157.764  (4.277) | 10.934  (1.687) | 72.624  (2.25) | 4.834  (0.036) | 5.877  (0.116) | 0.488  (0.004) | 45.432  (5.45) | 2.712  (0.813) | 17.384  (0.766) |
|  | Submaximum load lifting |  | 174.845  (4.724) | 19.056  (2.596) | 73.763  (2.575) | 4.849  (0.022) | 5.853  (0.074) | 0.488  (0.004) | 40.246  (6.357) | 2.236  (0.787) | 15.943  (0.804) |
| **Mid- upstroke** | Visual mask |  | 87.858  (5.082) | 0.208  (6.431) | 56.704  (4.135) | 4.709  (0.055) | 5.474  (0.133) | 0.479  (0.004) | 68.806  (5.333) | 4.228  (1.004) | 14.469  (0.673) |
|  | Typical flight |  | 90.357  (5.863) | 2.302  (2.41) | 40.584  (2.397) | 4.743  (0.046) | 5.48  (0.116) | 0.479  (0.003) | 74.342  (3.72) | 4.755  (0.735) | 12.845  (0.449) |
|  | Submaximum load lifting |  | 95.914  (8.45) | 5.794  (2.489) | 33.795  (2.888) | 4.755  (0.021) | 5.455  (0.053) | 0.479  (0.003) | 76.891  (1.396) | 4.622  (0.365) | 12.438  (0.655) |

**Movie 1** Example wing configurations during typical hoverfeeding, while hoverfeeding in front of a visual mask, and while hoverfeeding together with submaximum load lifting. Digitised landmarks are colour coded as in Figure 3.

**Movie 2** Example wing reconstructions during typical hoverfeeding, while hoverfeeding in front of a visual mask, and while hoverfeeding together with submaximum load lifting. *Left* Digitised landmarks. *Center* Wing surface reconstructed by joining points along the leading and trailing edges. *Right* Wing surface reconstructed by fitting through interior points on the wing.
